# WY14643 Increases Herpesvirus Replication Independently of PPARα Expression and Inhibits IFNβ Production

**DOI:** 10.1101/2021.10.29.466551

**Authors:** Lili Tao, Phillip Dryden, Alexandria Lowe, Guoxun Wang, Igor Dozmorov, Tyron Chang, Tiffany A. Reese

## Abstract

Peroxisome proliferator activated receptor (PPAR) agonists are commonly used to treat metabolic disorders in humans because they regulate fatty acid oxidation and cholesterol metabolism. In addition to their roles in controlling metabolism, PPAR agonists also regulate inflammation and are immunosuppressive in models of autoimmunity. We aimed to test whether activation of PPARα with clinically relevant ligands could impact herpesvirus infection using the model strain murine gammaherpesvirus-68. We found that PPARα agonists WY14643 and fenofibrate increased herpesvirus replication *in vitro*. *In vivo*, WY14643 increased viral replication and caused lethality in mice. Unexpectedly, these effects proved independent of PPARα. Investigating the mechanism of action for WY14643, we found that it suppresses production of type I interferon by inhibiting stimulator of interferon (STING), which lies downstream of the cytoplasmic DNA sensor cGAS. Thus, WY14643 regulates interferon downstream of cytoplasmic DNA recognition and increases herpesvirus replication in a PPARα-independent manner. Taken together, our data indicate that caution should be employed when using PPARα agonists in immuno-metabolic studies, as they can have off-target effects on viral replication.

**Importance:** PPAR agonists are used clinically to treat both metabolic and inflammatory disorders. Because viruses are known to rewire host metabolism to their own benefit, the intersection of immunity, metabolism, and virology is an important research area. Our article is an important contribution to this field because for two reasons. First, it shows a role for PPARα agonists in altering virus detection by cells. Second, it shows that PPARα agonists can affect virus replication in a manner unrelated to their expected genetic function. This knowledge is valuable for anyone seeking to use PPARα agonists as a research tool.

## Introduction

Viruses manipulate host cellular machinery, including metabolic pathways, to aid their own replication or to suppress host immune defenses. For instance, many human herpesviruses rewire glycolysis, glutaminolysis, and fatty acid synthesis to promote nucleotide synthesis and lipogenesis, allowing the host cell to sustain the high energy demands during viral infection and replication (1–5). Additionally, recent studies show that metabolic products inhibit type I interferon production downstream of both RNA and DNA sensing pathways (6, 7), suggesting the possibility that metabolically-active pharmaceuticals might affect virus infection.

To determine whether and how such compounds might impact virus infection and the immune system, we initially focused on the possible interactions of herpesvirus infection with compounds that target peroxisome proliferator activated receptors (PPARs). We chose to study herpesvirus/PPAR interactions because activation of all three PPAR isoforms—PPARα, PPARβ/δ, and PPARγ—profoundly impacts cellular metabolism and inflammatory signaling cascades (6). PPARα activation induces fatty acid oxidation and lowers intracellular lipid levels. PPARγ receptors are primarily expressed in adipose tissue and control lipogenesis and insulin sensitivity. PPARβ/δ receptors also orchestrate fatty acid oxidation, particularly in muscle (6). Consistent with their roles in cellular fatty acid metabolism, activation of PPARs by agonists such as fibrates (PPARα) and thiazolidinediones (PPARγ) has demonstrated clinically efficacy in treating metabolic disorders in humans (7–9). Importantly, fatty acid metabolism has been shown to regulate host-virus interaction and contribute to determining the outcome of infection (10, 11).

Moreover, PPARs regulate inflammation, and PPAR agonists are used clinically to reduce inflammation in atherosclerosis, diabetes, neurodegenerative diseases, and autoimmune diseases (8). Working through diverse pathways, PPAR agonists repress NF-κB and AP-1 DNA binding, regulate nitric oxide production, inhibit dendritic cell maturation, reduce cytokine expression by effector T cells, and inhibit leukocyte recruitment to sites of inflammation, all of which are known key regulators of cellular antiviral response (12, 13). Collectively, this evidence highlights the potential roles of PPARs in shaping immune responses against DNA viruses.

Conversely, it is known that herpesviruses manipulate host cell metabolism during infection to promote viral replication and chronic infection (14, 15), including through the induction of peroxisomes (16, 17). A recent report found that HCMV induces peroxisome biogenesis to enhance plasmalogen synthesis, which is required for efficient HCMV envelopment (17). Herpesviruses also encode viral proteins that target peroxisomes, suggesting that modulation of peroxisomal function is important for these viruses (18–20).

Despite their immunoregulatory functions, our understanding of the effects of PPAR agonists on infectious disease outcomes remains incomplete. There are contradictory reports suggesting that synthetic agonists or dietary lipids improve or impair resistance to pathogen challenge, and the molecular mechanisms of PPAR-mediated immunoregulation during infection remain elusive (21–25).

Thus, we sought to test the hypothesis that PPAR agonists could alter host-herpesvirus interactions and identify the underlying molecular mechanisms of such an effect. We evaluated agonists of PPARα, PPARβ/δ, and PPARγ and found that the compounds WY14643 and fenofibrate (agonists of PPARα) produce strong proviral effects. Treatment with these compounds dramatically increased the replication of herpesviruses in two different taxa by antagonizing the cytosolic DNA sensing pathway. Consistent with our hypothesis, this effect initially seemed dependent on PPARα expression. However, anomalous data led us to carefully control for mouse genetic background and microbiome and revealed that these proviral effects occur independently of PPARα.

## Results

### *PPARα agonists promote herpesvirus replication* in vitro

To examine the effects of PPAR activation on DNA virus infection, we used murine gammaherpesvirus-68 (MHV68) as our model. MHV68 readily infects mice and undergoes phases of infection similar to human gamma-herpesviruses such as Kaposi’s sarcoma associated herpesvirus (KSHV) and Epstein Barr virus (EBV) (26, 27). The replication of MHV68 was measured in bone marrow-derived macrophages (BMDMs), as this virus infects and replicates in macrophages as well as B cells and dendritic cells *in vivo* (26, 28)

The effects of PPAR activation on MHV68 replication were examined by treating BMDMs with agonists for PPARα (fenofibrate or WY14643), PPARβ/δ (GW501516), or PPARγ (rosiglitazone) prior to infection. After infection, we quantified viral replication with flow cytometry, staining for the expression of lytic viral proteins on the surfaces of infected cells (29). We found that PPARα agonists fenofibrate and WY14643 both increased expression of lytic viral proteins on infected macrophages (Fig. 1A). However, GW501516 and rosiglitazone had no effect on MHV68 replication, indicating that PPARβ/δ and PPARγ agonists do not regulate MHV68 replication (Fig. 1A). We confirmed these effects of WY14643 and fenofibrate with viral growth curves, which showed increased MHV68 replication in macrophages treated with fenofibrate or WY14643 compared to untreated cells (Fig. 1B, C). This was true at both high and low multiplicity of infection (MOI). Thus, treatment with PPARα agonists increases MHV68 replication.

**Figure 1.**
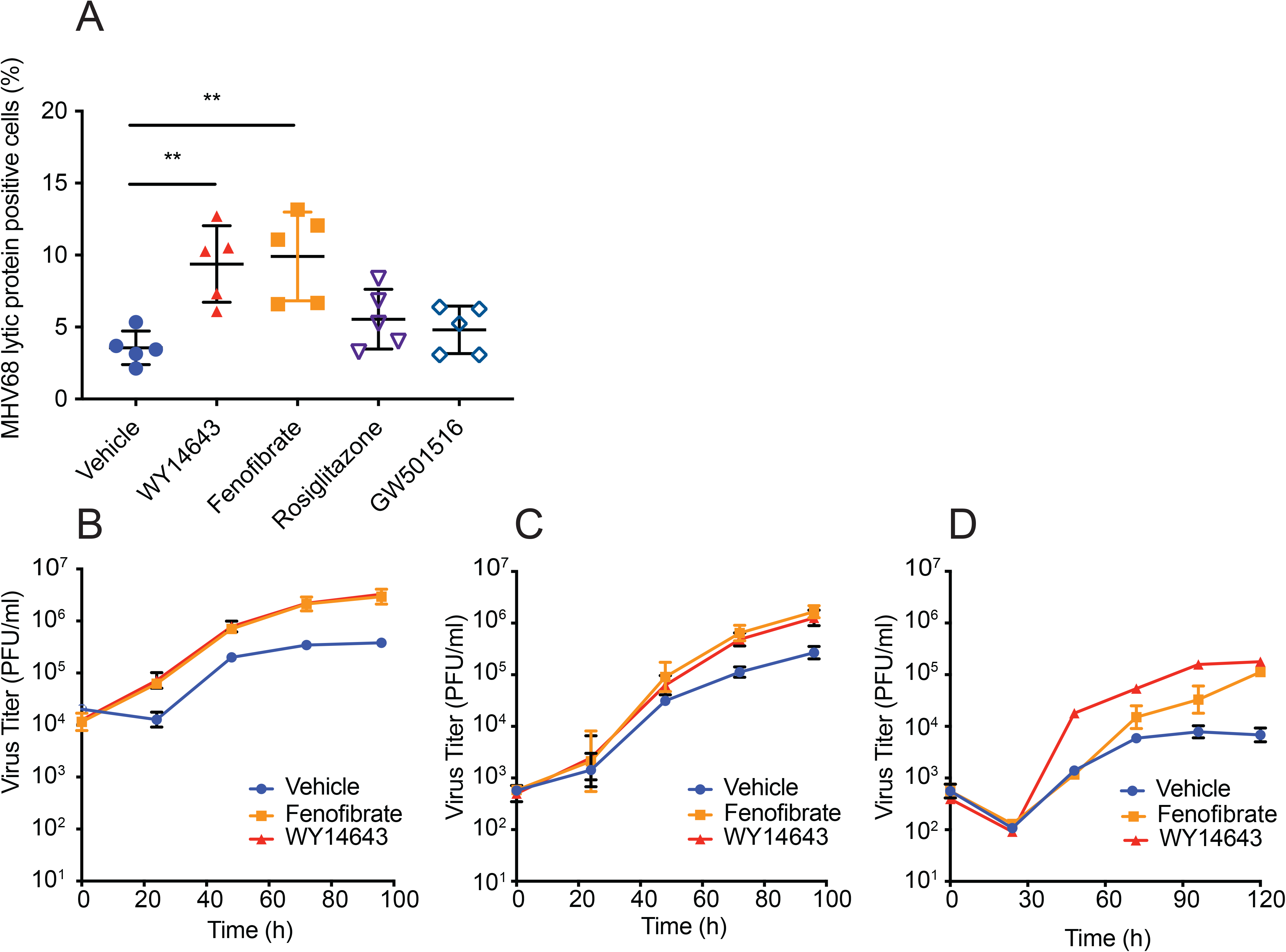
PPARα agonist treatment increases herpesvirus replication *in vitro*. A. BMDMs from C57BL6/J mice were pretreated for 16 hours with WY14643 (200 μM), fenofibrate (50 μM), rosiglitazone (1 μM), or GW501516 (100 nM). Cells were infected with MHV68 at MOI=5. The percentage of cells expressing MHV68 lytic proteins was measured with flow cytometry using a polyclonal antibody 24 hours after infection. The data represent the mean ± SD from 5 independent experiments. **p<0.01 by one-way ANOVA with Tukey’s multiple comparisons. B-C. Shown are growth curves of MHV68 in macrophages isolated from C57BL/6J mice after pretreatment with vehicle control (DMSO), WY14643 or fenofibrate. Cells were infected with MHV68 at MOI=5 (B) or MOI=0.1 (C). Virus was quantitated by plaque assay on 3T12 cells. The data represent the mean ± SD from 3 independent experiments. D. Shown are growth curves of MCMV in BMDMs isolated from C57BL/6J mice. After pretreatment with vehicle, WY14643, or fenofibrate, cells were infected with MCMV at MOI=1. The data represent the mean ± SD from 2 independent experiments.

To determine if PPARα agonists affect replication of other herpesviruses, we tested whether WY14643 or fenofibrate would increase replication of murine cytomegalovirus (MCMV), a betaherpesvirus, in macrophages. We found that replication of MCMV was also increased by these treatments (Fig. 1D), suggesting that PPARα agonists can modulate herpesvirus replication across taxa.

### WY14643 increases virus replication and lethality in mice independently of expression of PPARα

We next determined if the effects of PPARα agonists WY14643 and fenofibrate depend on expression of PPARα. Initially, we performed experiments in BMDMs isolated from *Ppara^−/−^* mice obtained from Jackson Laboratories (here denoted “*Ppara^−/−^/J*”), which we maintained with a knockout-to-knockout breeding scheme. In these experiments, the effects of fenofibrate and WY14643 were abolished in the *Ppara^−/−^/J* BMDMs (Fig. 2A). This suggested that their proviral activity involves their canonical role as PPARα agonists. However, we noted an increase in virus replication in vehicle-treated *Ppara^−/−^/J* macrophages relative to control macrophages from C57BL6/J mice (Fig. 2A).

**Figure 2.**
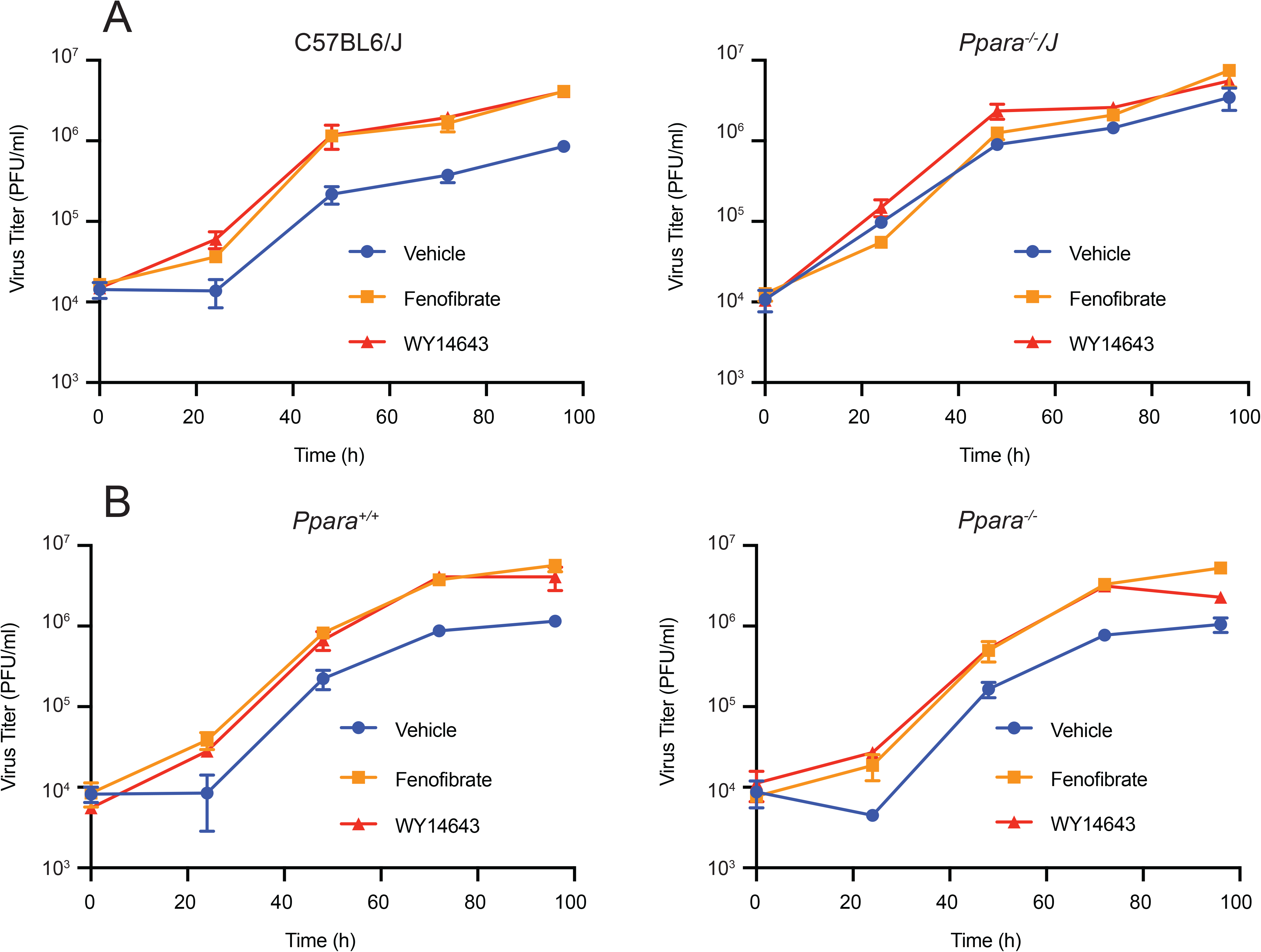
Littermate controls reveal that WY14643 and fenofibrate increase viral replication independently of PPARα *in vitro*. A. Shown are growth curves of MHV68 in BMDMs isolated from either C57BL6/J or *Ppara^−/−^/J* mice. Cells were pretreated with vehicle control, WY14643, or fenofibrate and then infected (MOI=5). Viral growth was quantitated by plaque assay on 3T12 cells. Data represent the mean ± SD of 2 independent experiments. B. Shown are growth curves of MHV68 in BMDMs isolated from littermate-controlled *Ppara^+/+^ or Ppara^−/−^* mice. Cells were pretreated with vehicle control, WY14643, or fenofibrate and then infected (MOI=5). Viral growth was quantitated by plaque assay on 3T12 cells. Data represent the mean ± SD of 2 independent experiments.

Questioning whether this was due to differences in mouse genetic background or microbiome differences between knockout and C57BL6/J mice, we generated littermate-controlled mice by crossing C57BL6/J mice with *Ppara^−/−^/J* mice to obtain heterozygous offspring. These heterozygous mice were interbred to generate both knockout (“*Ppara^−/−^*”) and wild type (“*Ppara^+/+^*”) animals which were used for BMDM generation. When we compared viral replication in these littermate-controlled macrophages, we observed no baseline difference in virus replication between the genotypes in the vehicle group (Fig. 2B). However, the effects of fenofibrate and WY14643 remained intact in the littermate-controlled knockout BMDMs; agonist treatment was associated with increased virus replication in PPARα-deficient cells just as in wild type cells (Fig. 2B). This suggests that fenofibrate and WY14643 increase virus replication independently of PPARα expression.

We confirmed these results with *in vivo* experiments, testing whether WY14643 would increase viral replication in mice. First, we used C57BL6/J and *Ppara^−/−^/J* mice to test the effects of *in vitro* treatment with WY14643. Mice were injected with WY14643 or a vehicle control for 7 days, starting 3 days prior to infection and continuing for 4 days after infection (Fig. 3A). Using luciferase-tagged MHV68 (MHV68-M3FL), we infected mice and imaged them over multiple days to measure acute virus replication (29, 30). We found that C57BL6/J mice treated with WY14643 had increased virus replication compared to vehicle-treated mice (Fig. 3B). This elevated virus replication was absent in *Ppara^−/−^/J* mice treated with WY14643 (Fig. 3D). Surprisingly, even though mice were infected with a dose of MHV68 that does not cause lethality in wild type mice, we observed that C57BL6/J mice treated with WY14643 succumbed to infection (Fig. 3C) at a frequency similar to mice deficient in the type I interferon receptor (31, 32). This lethality effect was not present in the knockout mice (Fig. 3C). As with our original *in vitro* data, these *in vivo* data suggest that WY14643 increases lethality and virus replication through its canonical function as an agonist of PPARα.

**Figure 3.**
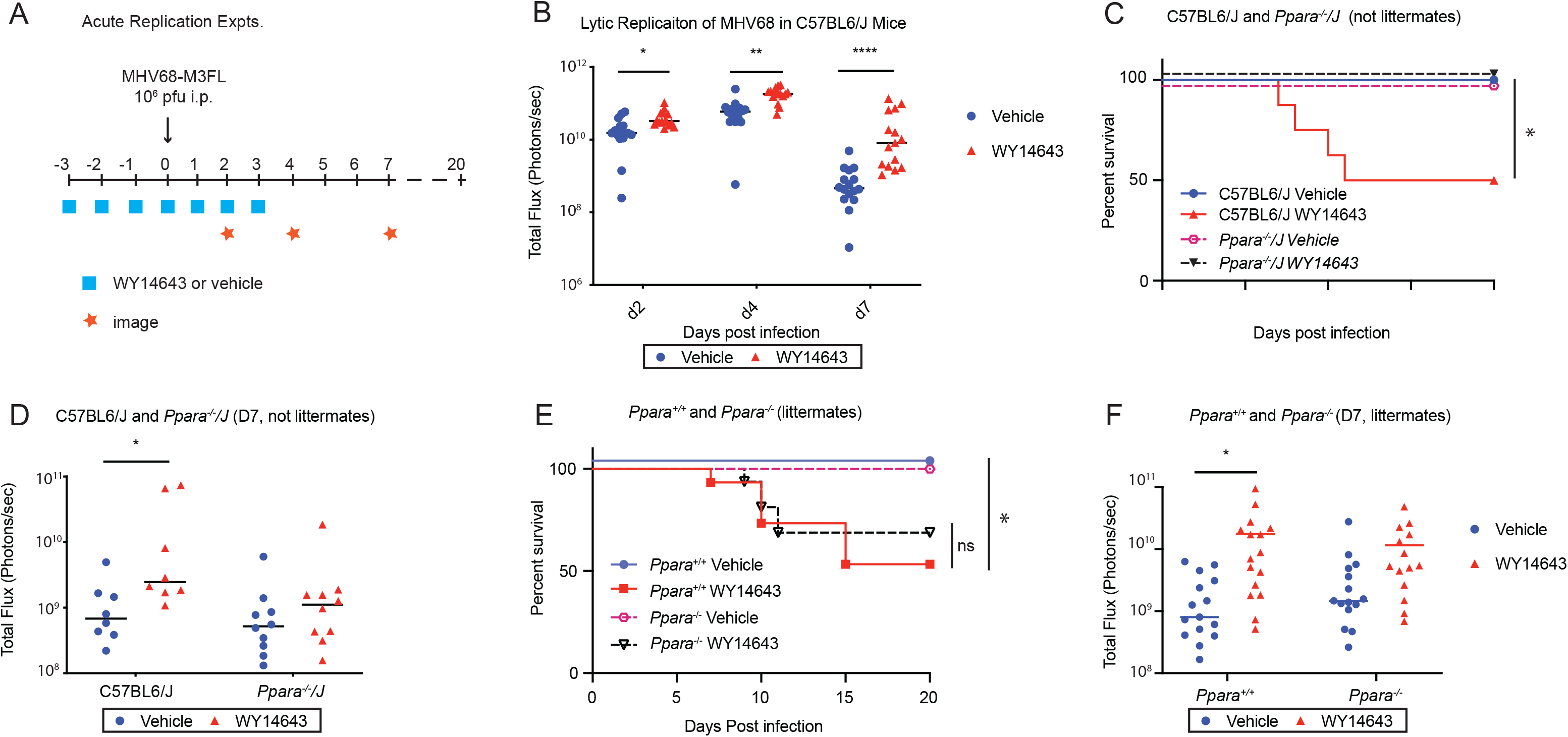
Littermate controls reveal that WY14643 increases viral replication and causes mortality independent of PPARα *in vivo*. A. Shown is a schematic prodedure for measuring acute replication of MHV68 in mice. Mice were injected intraperitoneally with either vehicle control (15% HS15 in normal saline) or WY14643 (100 mg drug in HS15 and normal saline per mouse kg) for 7 days, starting 3 days before infection. Mice were infected intraperitoneally with MHV68-M3FL at dose of 10^6^ PFU. Acute replication of virus was measured at d2, d4, and d7 after infection using an IVIS bioluminescence imager. Survival of mice was monitored until 20 days after infection. B. Shown is the acute replication of MHV68 over time in C57BL6/J mice measured by luminescent intensity. Mice were treated with vehicle or WY14643. The data shown are the pool of 3 independent experiments. C. Shown are the survival rates of mice infected with MHV68. C57BL6/J (n=16) or *Ppara^−/−^/J* n=20) mice were treated with either a vehicle control or WY14643 and infected with MHV68. Data are shown as the pool of 2 independent experiments. *p<0.95 by log-rank (Mantel-Cox) test. D. Shown is the acute replication of MHV68 in C57BL6/J or *Ppara^−/−^/J* mice treated with vehicle control or WY14643 7 days after infection as measured by luminescent intensity. Data are shown as the pool of 2 independent experiments. E. Shown are the survival rates of littermate-controlled mice infected with MHV68. *Ppara^+/+^* (N=30) and *Ppara^−/−^* (N=31) mice were treated with either vehicle control or WY14643, and infected with MHV68. Data are shown as the pool of 3 independent experiments. *p<0.95 by log-rank (Mantel-Cox) test. F. Shown is the acute replication of MHV68 in littermate-controlled *Ppara^+/+^* (n=30) or *Ppara^−/−^* (n=31) mice measured by luminescent intensity. The data are shown as the pool of 3 independent experiments. Data all shown as mean ± SD; ^*^p<0.05, ^**^p<0.01, ^***^p<0.001, ****p<0.0001 statistical analysis was conducted using one-way ANOVA, two-way repeated measures ANOVA tests, or with Sidak’s multiple comparison test.

However, after repeating these experiments with mice interbred in our colony, we no longer observed the PPARα-dependency of WY14643. Littermate-controlled knockout mice treated with WY14643 succumbed to virus infection at a similar rate to wild type littermate controls (Fig. 3E) and also displayed increased virus replication (Fig. 3F).

At first, our *in vitro* and *in vivo* data suggested that the proviral effects of fenofibrate and WY14643 depend on PPARα. However, this seems to be an artifact of different mouse genetic backgrounds or microbiome differences between the C57BL6/J and knockout colony; when we used littermate controls, this dependency vanished. This indicates that fenofibrate and WY14643 increase MHV68 replication and animal mortality independently of their canonical roles as agonists of PPARα.

### WY14643 suppresses the interferon response by reducing type I IFN production

We next sought to identify how PPARα agonists alter the immune response to virus infection. To do this, we examined the broad transcriptional response to WY14643 treatment and MHV68 infection of macrophages with RNA sequencing analysis. Four groups of BMDMs were analyzed: uninfected with vehicle treatment, infected with vehicle treatment, uninfected with WY14643 treatment, and infected with WY14643 treatment. Six hours after infection, BMDMs were collected and prepared for RNA sequencing.

Pathway analysis of differentially expressed genes revealed that virus infection increased expression of interferon responsive genes, genes involved in inflammation, and NFκB pathway genes (Fig. 4A, Supplemental Table 1). This effect was present in vehicle-treated and WY14643-treated cells but reduced in magnitude in the WY14643 group. This reduction in magnitude suggests that WY14643 attenuates the early antiviral response in infected cells. To confirm the sequencing results, we quantified the expression of a subset of interferon-stimulated genes with RT-qPCR. Measurement *of Isg20*, *Isg15*, and *Cxcl10* transcripts confirmed that, indeed, expression of these genes was reduced in both uninfected and infected macrophages treated with WY14643 (Fig. 4B).

**Figure 4.**
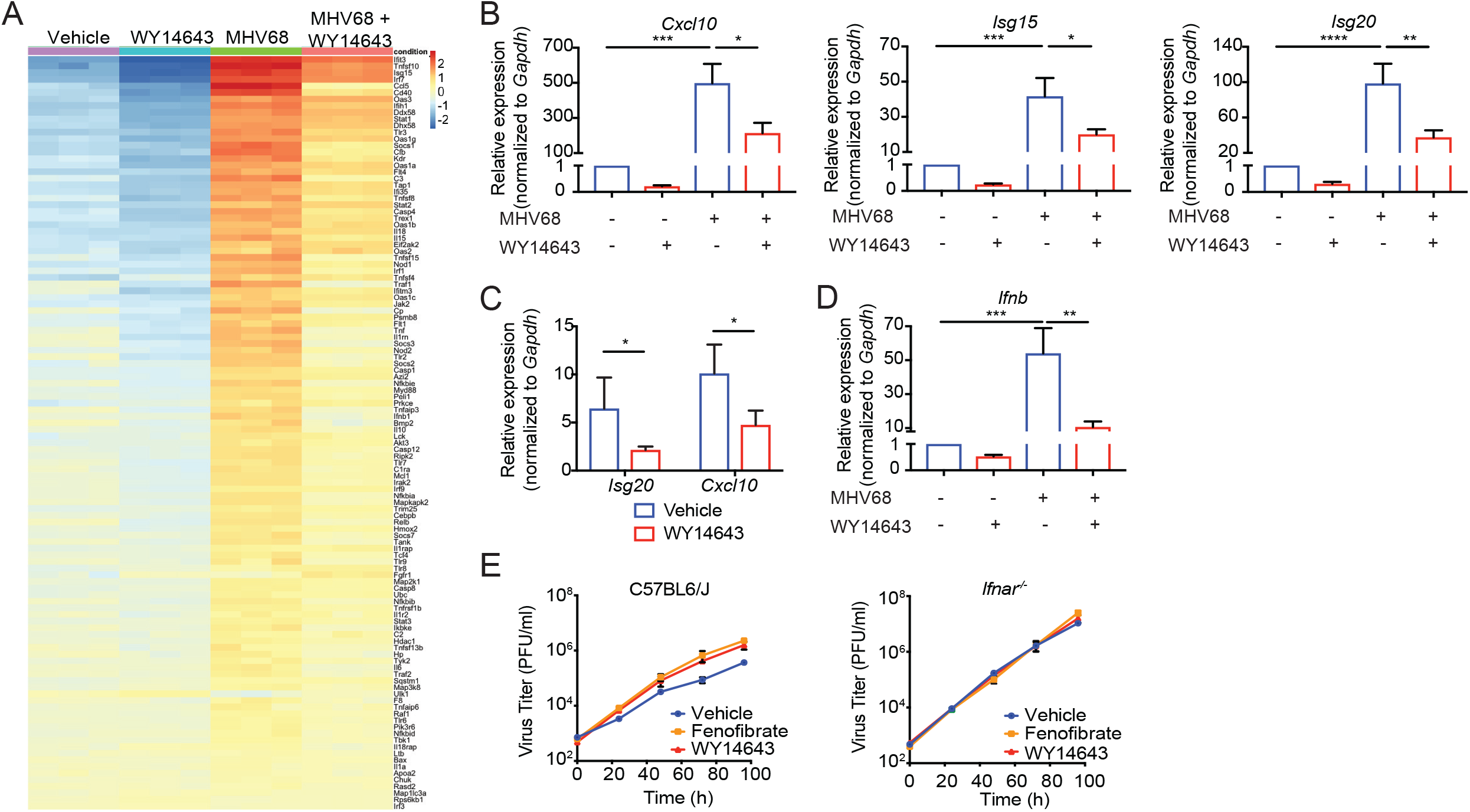
PPARα stimulation suppresses type I IFN. A. Macrophages were pretreated with a vehicle control or WY14643 for 16 hours prior to infection with MHV68 (MOI=5). 6 hours after infection, RNA was isolated and prepared for RNA sequencing. B. Macrophages were pretreated with vehicle or WY14643 for 16 hours prior to infection with MHV68. Then, qRT-PCR was performed to quantify transcripts of of *Cxcl10* (n=5), *Isg20* (n=7), and *Isg15* (n=6) before and 6 hours after MHV68 infection. Relative expression of these genes is shown normalized to *Gapdh*. C. Peritoneal exudate cells were collected from vehicle or WY14643-treated C57BL6/J mice (n=4) 2 days after infection with MHV68. Relative expression of *Isg20* and *Cxcl10* was quantitated and normalized to *Gapdh*. Data represent 2 independent experiments. D. Macrophages (n=8) were pretreated with vehicle or WY14643 for 16 hours prior to infection with MHV68. 6 hours after infection, qRT-PCR was performed to quantify *Ifnb*. The data represent relative expression normalized to *Gapdh*. E. Macrophages from C57BL6/J mice or *Ifnar^−/−^* mice were treated with vehicle, fenofibrate or WY14643 for 16 hours. Cells were infected with MHV68 at MOI=0.1 and viral growth was quantitated by plaque assay on 3T12 cells. The data shown the mean ± SD from 3 independent experiments. Data all shown as mean ± SD; ^*^p<0.05, ^**^p<0.01, ^***^p<0.001, ****p<0.0001 statistical analysis was conducted using one-way ANOVA with Tukey’s post-test.

We also tested these results *in vivo*, questioning whether mice treated with WY14643 and infected with MHV68 would display reduced expression of interferon-induced genes. To measure this, we analyzed interferon-stimulated gene expression in peritoneal cells from vehicle-treated and agonist-treated infected mice. We found that expression of *Isg20* and *Cxcl10* was decreased in agonist-treated mice (Fig. 4C), consistent with our *in vitro* data.

These data indicate that WY14643 acts through the type I interferon pathway, so we hypothesized that agonist treatment might reduce interferon production. To test this, we quantified *Ifnb* expression in uninfected and infected BMDMs. MHV68 infection induced *Ifnb* transcription as expected, but WY14643 treatment attenuated this effect (Fig. 4D). Because the type I interferon response is essential to control viral infection and WY14643 suppresses interferon-stimulated genes, we tested whether the effects of WY14643 acts through the type I interferon receptor, interferon alpha/beta receptor (IFNAR). We examined viral growth in wild type and *Ifnar^−/−^* macrophages with and without agonist treatment. As expected, virus grew substantially more in *Ifnar^−/−^* cells compared with wild type cells. WY14643 did not further increase virus replication in the knockout cells (Fig. 4E). Taken together, these data indicate that treatment with WY14643 during acute MHV68 infection suppresses the interferon response by reducing type I interferon production.

### WY14643 antagonizes interferon induction through the cytoplasmic DNA sensing pathway

Previously published work established that mice deficient in IFNAR have increased viral replication and enhanced susceptibility to MHV68 (31). Therefore, the increased lethality we observed in WY14643-treated mice was likely due to the suppression of type I interferon that we observed. Questioning the cause of this suppression, we hypothesized that WY14643 might act through the cytosolic DNA-sensing pathway. A DNA virus, MHV68 induces interferon production via cytosolic DNA sensing in the cyclic GMP-AMP synthase/stimulator of interferon gene (cGAS/STING) pathway. After detecting cytosolic DNA, cGAS produces the second messenger cGAMP, which activates STING. STING phosphorylates TBK1 and IRF3, leading to transcription of IFNβ. Therefore, we hypothesized that WY14643 could antagonize the early induction of interferon downstream cytoplasmic DNA recognition.

To test if suppression of IFNβ depends on the cGAS/STING pathway, we compared MHV68 replication in macrophages from *Sting^−/−^* and C57BL6/J mice that were treated either with WY14643 or a vehicle control. As expected, viral replication increased in the *Sting^−/−^*-vehicle group (Fig. 5A). However, *Sting^−/−^* macrophages treated with WY14643 did not display any further increase in virus replication (Fig. 5A). This suggests that the effects of WY14643 depend on STING expression. Although previous studies suggest that PPARs trans-repress NF-κB signaling and block transcription of cytokine genes (33), our result suggests that WY14643 can inhibit interferon by blocking the upstream signaling pathway.

**Figure 5.**
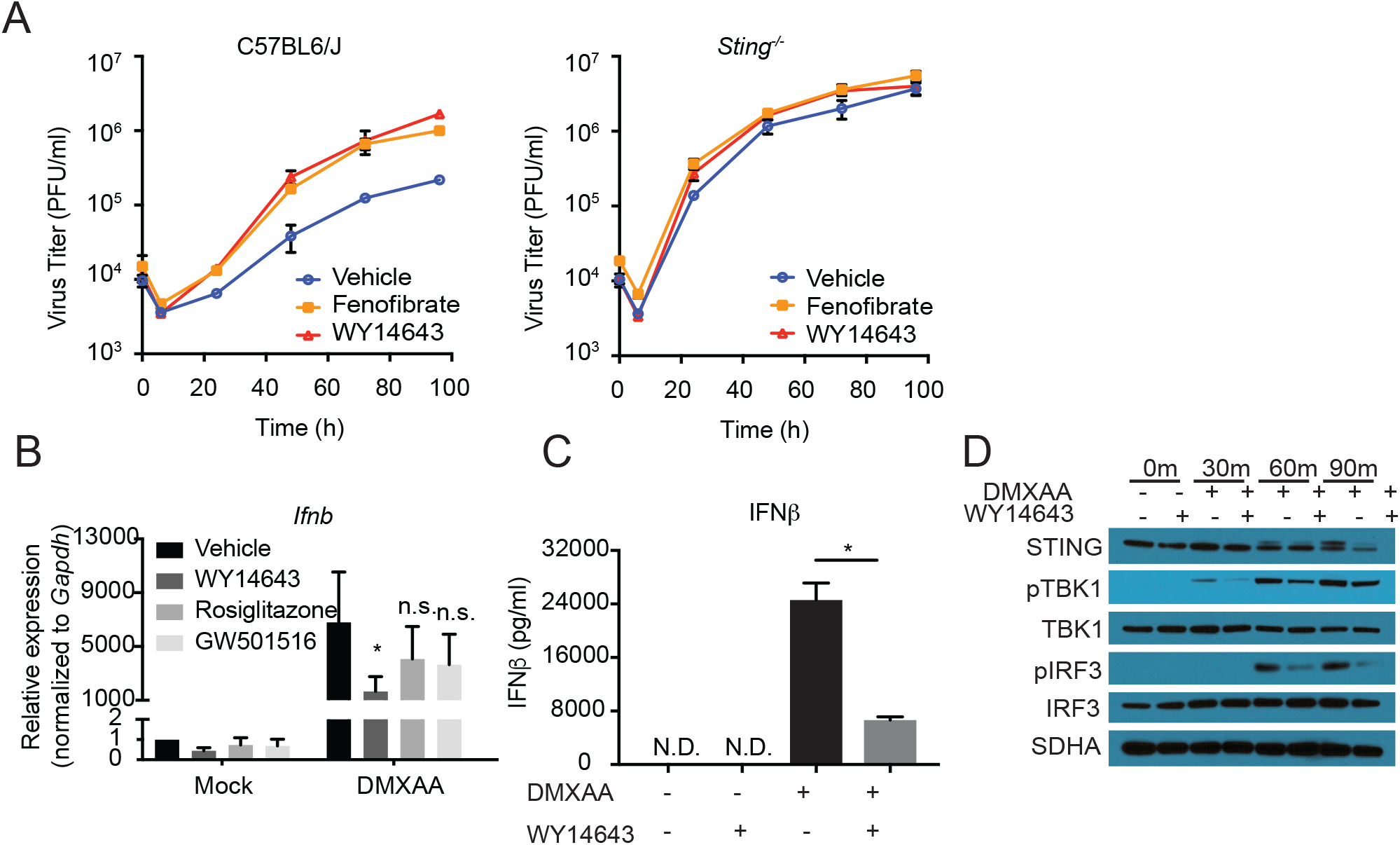
WY14643 and fenofibrate decrease the interferon response via the STING pathway. A. BMDMs from C57BL6/J or *Sting*^−/−^ mice were pretreated with vehicle or WY14643 for 16 hours prior to infection with MHV68, MOI=5. Virus growth was quantitated by plaque assay. The data shown the mean ± SD from 3 independent experiments. B. BMDMs from C57BL6/J mice were treated with a vehicle control, WY14643, fenofibrate, rosiglitazone, or GW501516 for 16 hours. Cells were then treated with DMXAA. 2 hours after treatment, qRT-PCR was performed with lysates and *Ifnb* expression was quantitated. The data was normalized to *Gapdh,* shown is the mean ± SD; ^*^p<0.05, n.s. (not significant), relative to DMXAA/vehicle treated. C. BMDMs were pretreated with vehicle or WY14643 for 16 hours prior to treatment with DMXAA. Concentration of IFNβ in culture medium before and 24 hours after transfection was determined by ELISA assay. The data show the mean ± SEM from 3 independent experiments. D. Macrophages were pretreated with the vehicle control or WY14643 for 16 hours. Shown is a representative western blot of proteins involved in STING signaling before transfection and 30 mins, 60 mins, 90 mins after transfection with DMXAA. Data all shown as mean ± SD; ^*^p<0.05, ^**^p<0.01, ^***^p<0.001,, ****p<0.0001 statistical analysis was conducted using one-way ANOVA test.

Because WY14643 requires STING to induce its effects, we hypothesized that WY14643 would suppress interferon production upon cytoplasmic DNA stimulation. To test this, we stimulated BMDMs with DMXAA, a murine agonist of STING, after applying vehicle or WY14643 treatment. As expected, DMXAA treatment resulted in increased *Ifnb* gene expression (Fig. 5B). Consistent with our hypothesis, this effect was significantly diminished in cells treated with WY14643, but not in those treated with rosiglitazone and GW501516 (Fig. 5B). We also measured IFNβ protein levels in macrophage supernatants and found that WY14643 treatment reduced IFNβ production after DMXAA stimulation (Fig. 5C). To determine if WY14643 inhibited STING signaling, we examined phosphorylation of TBK1 and the transcription factor IRF3. We found that WY14643 suppressed DMXAA-induced phosphorylation of both proteins (Fig. 5D). Therefore, we concluded that WY14643 suppresses IFNβ production downstream of cytoplasmic DNA sensing. Together with the data in *Sting^−/−^* macrophages, these data suggest that WY14643 antagonizes production of IFNβ by altering the ability of STING to activate TBK1 and IRF3.

## Discussion

We determined that two agonists of PPARα, WY14643 and fenofibrate, increase herpesvirus replication independently of expression of PPARα. In macrophages, WY14643 and fenofibrate both increased replication of both a gamma- and beta-herpesviruses and suppressed the induction of IFNβ. *In vivo*, WY14643 caused lethality in infected mice along with increased virus replication, a strong effect given that MHV68 is rarely fatal. We determined that the suppression of IFNβ was STING-dependent, suggesting that WY14643 regulates type I interferon production through the cytosolic DNA sensing pathway. Importantly, the effects of WY14643 on herpesvirus infection are independent of PPARα.

Our findings leave many questions open regarding the role of PPAR activation and virus infection. Although the phenotype we observed is independent of PPARα, our data do not rule out any possible PPAR-dependent interactions between PPAR agonists and virus infection. Such interactions remain worth investigating. One such mechanism is the upregulation of negative regulators of inflammation such as IκB and the soluble IL-1 receptor antagonist (34). A second mechanism is by regulating inflammatory gene expression directly. PPARs decrease NFκB and AP-1 activities through transrepression, which stabilizes corepressor complexes on inflammatory gene promoters, such as nitric oxide synthetase, IL-1β, and IL-12 (33, 35). A third potential mechanism is by inducing peroxisomal metabolism, which may impact antiviral signaling pathways. PPARs, as their name implies, increase peroxisomal metabolism. Peroxisomes synthesize phospholipids and bile acids and oxidize very long, branched chain, and polyunsaturated fatty acids. They also produce significant amounts of hydrogen peroxide and other reactive oxygen species. In addition, fibrates and WY14643 induce ROS (36–40). As we recently reported, the induction of ROS in MHV68-infected cells or cells stimulated via the cGAS/STING pathway reduces production of IFNβ by impairing STING polymerization (41).

Each of these pathways remain likely targets for investigation, though our results indicate that researchers should be mindful of PPAR-independent immune modulation caused by PPAR agonists. This is especially true because some of our findings resembled the phenotype we might predict based on a PPAR-dependent mechanism. For instance, our RNA sequencing data showed comparative downregulation of NF-κB-pathway genes, interferon response genes, and inflammatory genes in WY14643-treated cells. Additionally, our experiments identified that WY14643 increases herpesvirus infection by antagonizing type I interferon production downstream of STING. All of these data would be consistent with a PPARα-dependent mechanism for WY14643, though careful controls revealed that WY14643 acts independently of PPARα.

Separating PPARα-dependent and independent effects may be complicated by genetic modifiers and microbiome differences in mouse models, as was the case in our experiments. Initially, we observed a PPARα-dependent phenotype when comparing *Ppara^−/−^/J* and C57BL6/J macrophages. The effects of WY14643 and fenofibrate on virus replication were abolished in *Ppara^−/−^/J* cells, suggesting that their mechanism is through their canonical role as PPARα agonists. However, MHV68 replication was unexpectedly high in vehicle-treated *Ppara^−/−^/J* macrophages compared with the control, leading us to question the validity of the comparison. This anomalous virus replication, along with the illusion of PPARα-dependency, disappeared when we generated and used littermate-controlled *Ppara+/+* and *Ppara^−/−^* mice. There are several possible reasons for this. For one, the Jackson Laboratory does report that the *Ppara^−/−^/J* mice have 3 single-nucleotide polymorphism markers that are still of the 129S4/SvJae allele-type, which could contribute to differences in immune response and virus replication. Additionally, microbiome differences between our C57BL6/J and *Ppara^−/−^/J* colonies, which were normalized by interbreeding the two, could have played a role. The exact causes of the false PPARα-dependent phenotype remain unclear, but highlight the importance of using littermate controls when trying to establish genetic dependency.

We set out to examine the role of PPARα in virus infection using the pharmacological agonist WY14643, as well as fenofibrate. We found that these compounds alter the STING-dependent antiviral immune response. Additionally, we found that treatment with these compounds increases herpesvirus replication *in vitro.* WY14643 had a profound effect *in vivo*, causing increased replication and lethality in mice infected with MHV68, a virus that is not normally fatal. Importantly, these effects on virus replication are independent of PPARα expression. These results are similar to other reports describing the off-target effects of etomoxir, an inhibitor commonly used to block Cpt1a and fatty acid oxidation (42–44). Our data indicate that caution is warranted when interpreting pharmacological studies that do not establish genetic specificity.

## Acknowledgments

We thank members of the Reese labs for technical assistance, Nan Yan, David Mangelsdorf and Steven Kleiwer for reagents and expertise, Dr. Wenhan Zhu for help with data analysis. We also thank the UTSW Flow Cytometry core and the Genomic and Microarray core for technical assistance. The Reese lab is supported by the NIH (1R01AI130020-01A1 and 5U19AI142784), CPRIT (RP200118), the Pew Scholars Program.

## Author Contributions

L.T. and P.D. performed and analyzed experiments, with technical assistance from A.L, G.W and T.C. I.D. performed RNAseq analysis. T.A.R. directed the project and wrote the manuscript with the assistance of P.D.

## Material and Methods

### Animals

C57BL/6J, B6;129S4-*Pparα*^tm1Gonz^/J (45), and B6.129S2-*Ifnar1^tm1Agt^*/Mmjax (46), were purchased from The Jackson Laboratory. All mice were housed under specific pathogen-free, double-barrier facility at the University of Texas Southwestern Medical Center. Mice were fed autoclaved rodent feed and water. Mice were maintained and used under a protocol approved by UT Southwestern Medical Center Institutional Animal Care and Use Committee (IACUC).

### Chemicals

PPARα agonist WY14643 was purchased from Cayman Chemical. PPARα agonist fenofibrate, PPARβ agonist GW501516, and PPARγ agonist rosiglitazone were purchased from Sigma Aldrich.

### Cell culture

Bone marrow derived macrophages were differentiated in DMEM (Corning) with 10% FBS supplemented with 1% glutamine (Corning), 1% HEPES (Corning) and 10% CMG14 supernatant for 7 days (47). 3T12 cells were maintained in DMEM with 5% FBS supplemented with 1% glutamine and 1% HEPES.

### Generation of virus stocks

Murine γ-herpesvirus 68 (WUSM stain) was purchased from ATCC. Murine γ-herpesvirus 68-M3FL was generated as previously reported (30).

### Virus infection

Fully differentiated BMDMs were seeded on 24 well plates (1.5×10^5^ cells per well) or 6 well plate (10^6^ cells per well). Cells were pretreated with either vehicle control (0.1% DMSO) or agonists (Fenofibrate 50 μM, WY14643 200 μM, Rosiglitazone 1 μM, GW501516 100 nM) for 16 hours. The next days, macrophages were infected with MHV68 at multiplicity of infection (MOI)=5 or 0.1. For MCMV experiments, cells were infected at MOI=1. After an hour, cells were washed with PBS twice to remove unabsorbed viruses. Then, culture medium containing treatments was added to the wells. For growth curve, samples were collected at 0 hour, 24 hours, 48 hours, 72 hours and 96 hours after infection and were frozen at −80 °C. The titer of virus was determined by plaque assay in 3T12 cells. For flow cytometric analysis, cells were collected 24 hours after infection.

### Flow cytometry for MHV68 lytic proteins positive cells

To determine the percentage of cells that expressing lytic proteins of MHV68 infection, cells were harvested 24 hours after infection, and fixed with 2% formaldehyde, blocked with 10% mouse serum and 1% Fc block (CD16/32, BioLegend), then stained with polyclonal rabbit antibody to MHV68 (1:1000) (29, 48), followed by secondary goat anti-rabbit Alexa Fluor-647 (Invitrogen).

### Plaque assay

The concentration of virus was titered in 3T12 cells. The frozen samples containing viruses were thawed in an incubator. The samples were serial diluted, then added to a monolayer of 3T12 cells. After an hour of absorption, the cells were then covered with 1% methylcellulose. Plates were incubated at 37 °C for 7 days, and the plaques were stained with 0.1% crystal violet.

### RT-qPCR

BMDMs in 6 well plates were either infected with MHV68 at MOI=5 for 6 hours or treated with STING ligand DMXAA at 10 μg/ml for 2 hours. RNA was extracted using Qiagen RNeasy Mini Kit (Qiagen) and reverse transcribed into cDNA using SuperScript VILO cDNA Synthesis Kit (Thermo Fisher Scientific). Relative quantification of target genes was determined using PowerUp SYBR Green Master Mix (Thermo Fisher Scientific) in a QuantStudio 7 Flex real time PCR system. Primers used for amplifying target genes are listed in Table 2.

**Table 2.**
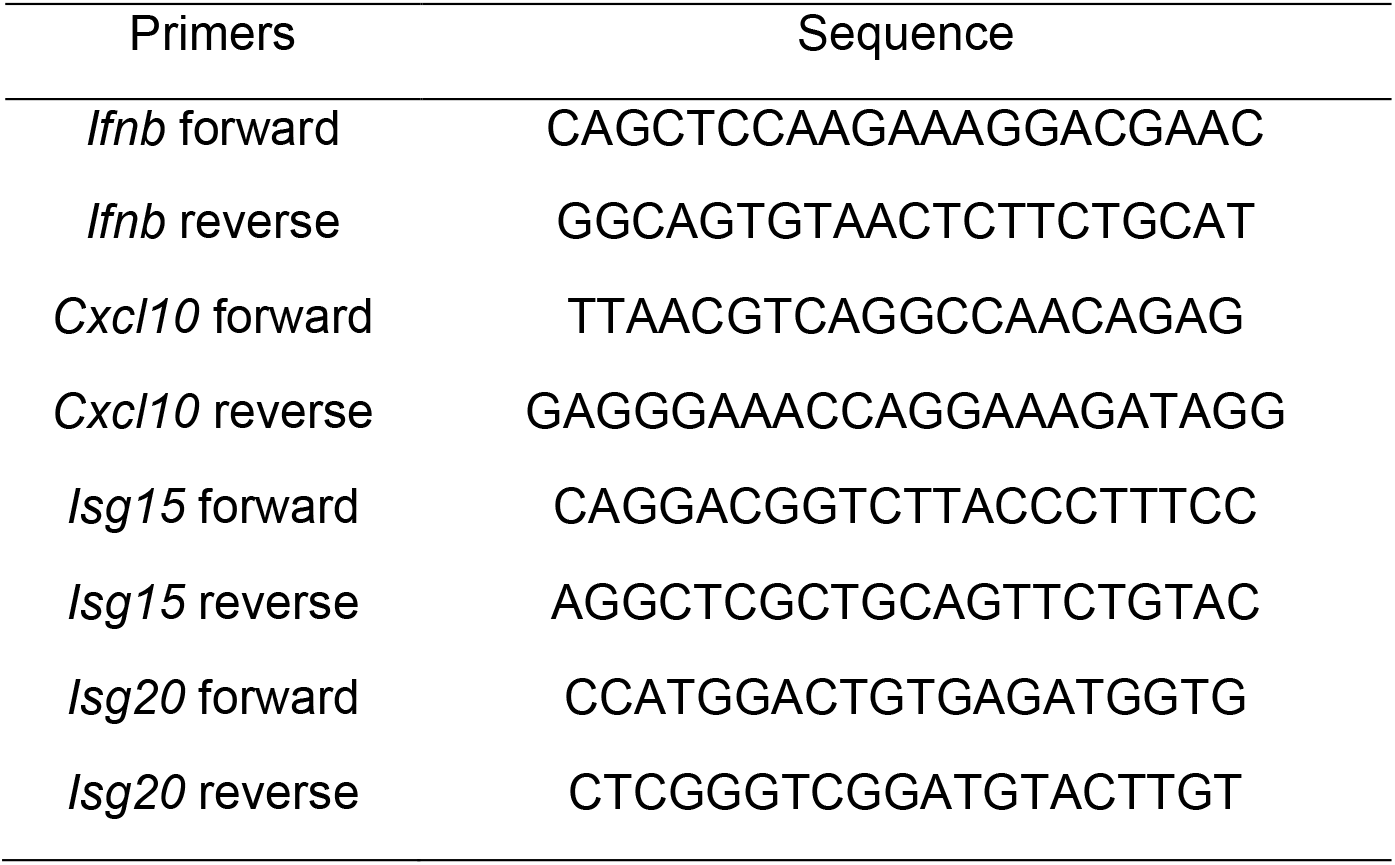
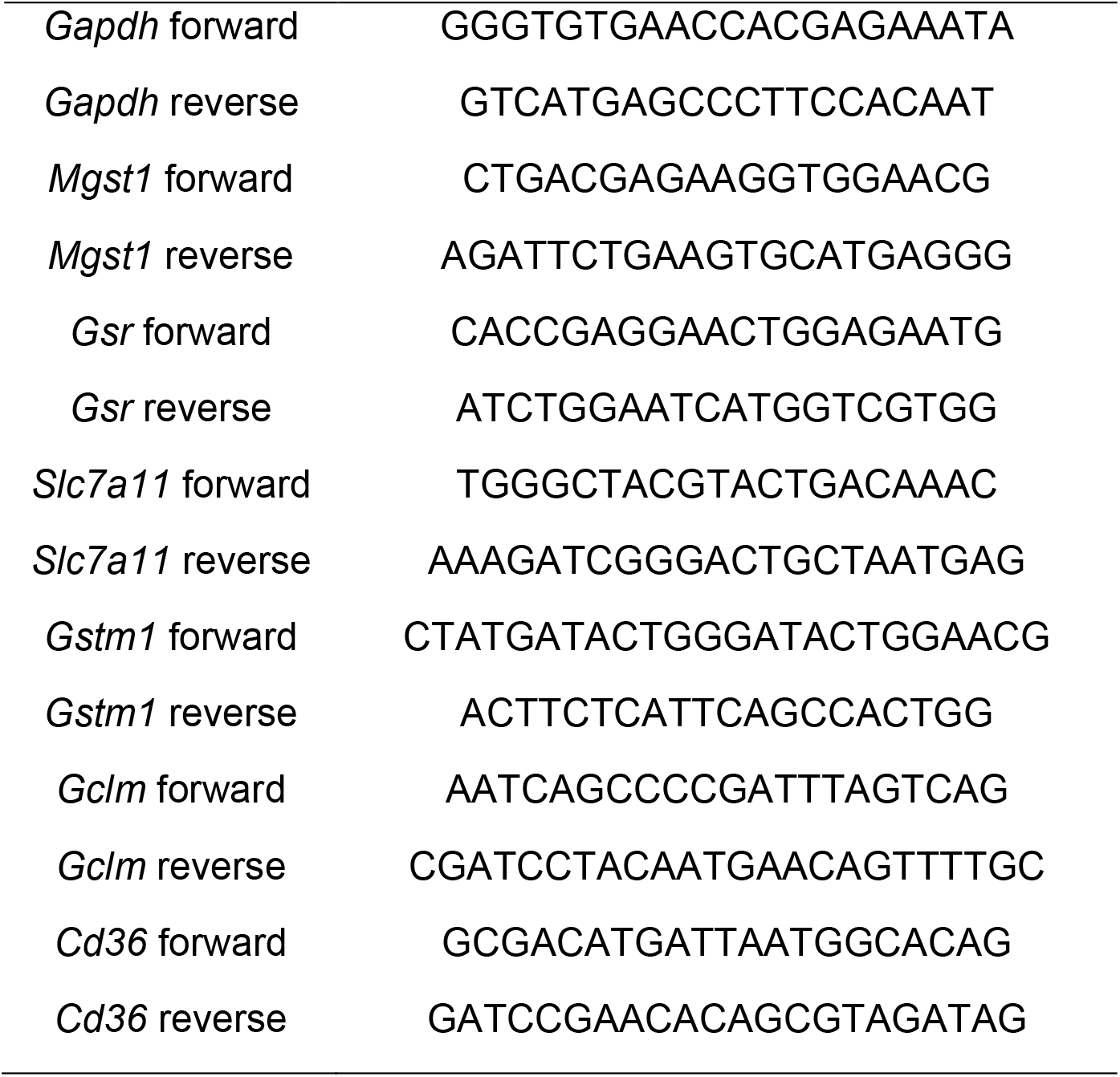
QPCR primers for target genes

### IFNβ ELISA Assay

BMDMs were seeded in 24 well plates. The next day, cells were treated with either vehicle control or WY14643 for 16 hours. Cells were then treated with 10 μg/ml DMXAA. Supernatant of cells were collected 24 hours after treatment and frozen at −80 °C. The concentration of IFNβ in the supernatant was determined with PBL IFN Beta ELISA Kit (PBL Assay Science).

### MHV68 acute replication in mice

Experiments were carried out using 8-12 weeks old mice under the protocol approved by IACUC. Mice were injected intraperitoneally with either vehicle control (15% HS15 in normal saline) or WY14643 (100 mg/kg) for 1 week starting from 3 days before virus infection. Mice were then infected with MHV68-M3FL at the dose of 10^6^ PFU through intraperitoneal route (29, 30). To quantify virus-encoded luciferase expression, mice were weighed and injected with 150 mg/kg of D-Luciferin (GOLDBIO) immediately prior to imaging using IVIS Lumina III In Vivo Imaging System (PerkinElmer). Total flux (photons/second) of the abdominal region was measured (exposure=1 sec., F/stop= 1.2, FOV=E, binning=medium, emissions filter=open) using Living Image software (PerkinElmer). Survival of the mice was recorded until 20 days after infection.

### RNA sequencing and data analysis

RNA samples were extracted with RNeasy Mini Kit (Qiagen) and RNA library was prepared using TruSeq Stranded mRNA Sample Preparation Kit (Illumina). Samples were sequenced on NextSeq 500/500 sequencer (Illumina) with SE-85. RNA sequencing data is normalized and analyzed based on the use of “internal standards” (49) that characterize some aspects of the system’s behavior, such as technical variability, as presented elsewhere (50, 51). The two-step normalization procedure and the Associative analysis functions are implemented in MATLAB (MathWorks, MA) and available from authors upon request. Functional analysis of identified genes was performed with Ingenuity Pathway Analysis (IPA; Ingenuity System). The sequencing data has been deposited in the European Nucleotide Archive under the accession number PRJEB33753.

#### QUANTIFICATION AND STATISTICAL ANALYSIS

All data are presented as mean ± SD. Statistical comparisons were performed using GraphPad Prism 7.0 software. Data were compared using unpaired two-tailed t test, one-way or two-way ANOVA. Statistical significance was set at p < 0.05. The numbers of independent replicates (n) are reported in the figure legends.

